# Commercial porcine gastric mucin contributes to variation in production of small molecule virulence factors by *Pseudomonas aeruginosa* when cultured in different formulations of artificial sputum medium

**DOI:** 10.1101/2021.01.25.428197

**Authors:** Rachel L. Neve, Brent D. Carrillo, Vanessa V. Phelan

## Abstract

Site-specific *in vitro* culture media are being developed to investigate microbial pathogenicity and ecology in nutrient environments that are more reflective of disease. For microbial ecology research in cystic fibrosis (CF), different artificial sputum media (ASM) formulations have been created to recapitulate the nutrient availability of the CF lung environment. However, these ASM formulations vary in concentration of amino acids, mucin, and other niche-specific compounds. Here, we measured the differential production of small molecule virulence factors by *Pseudomonas aeruginosa*, the predominant pathogen in CF pulmonary infections, cultured in nine different ASM formulations via liquid chromatography tandem mass spectrometry (LC-MS/MS) and molecular networking. We show that different ASM formulations lead to different phenotypes and metabolic profiles of *P. aeruginosa* and commercial porcine gastric mucin (PGM) contains a myriad of contaminants, including iron, which affect *P. aeruginosa* physiology.

**IMPORTANCE:** Different media formulations aiming to replicate *in vivo* infection environments contain different nutrients, which affects interpretation of experimental results. Inclusion of undefined components, such as commercial porcine gastric mucin (PGM), in an otherwise chemically defined medium can alter the nutrient content of the medium in unexpected ways and influence experimental outcomes.

---

*In vitro* cultivation systems are being developed and applied to investigate interactions between members of host-associated microbiota using native-like nutritional environments. These systems are beneficial to researchers because they enable higher resolution analysis of microbiome community structure and function by removing confounders associated with host cells, immune response, and variability in nutritional intake.^1, 2^ Foundational to the utility of *in vitro* culture systems to investigate the microbiota is the replication of host sites using physiologically relevant media. However, across laboratories, different variations of site-specific culture media are being used in experimental investigations. A representative example of multiple media formulations created and used to interrogate site-specific microbial interactions is artificial sputum media (ASM), a class of media aiming to replicate the nutritional environment encountered by the pulmonary microbiome of persons with cystic fibrosis (CF).^3–14^ ASM have been applied to investigate microbial physiology^3, 5, 15–17^, biofilm morphology^10, 18^, antibiotic susceptibility^19, 20^, and interspecies interactions in the context of CF^21–23^.

CF is a hereditary disease characterized by the accumulation of thick and sticky bronchial mucus in the lungs, which enables growth of opportunistic pathogens.^24^ Pulmonary mucus is a nutritionally distinct environmental niche comprised of mucin, DNA, amino acids, lipids, and micronutrients, including metals^25, 26^. Reflective of the variability of measurements of nutrient concentration in CF sputum and bronchoalveolar lavage fluid (BALF)^27, 28^, each ASM formulation differs in the concentration mucin, DNA, amino acids, lipids, and metals, affecting overall nutrient composition of the media.^25, 26^ Variations in ASM formulation are driven by literature precedent, direct measurement of nutrients from clinical samples, personal observations, and economic cost of materials.^3–14^ Importantly, disparate nutrient concentration and bioavailability between ASM formulations may affect microbial growth, metabolism, and virulence^3, 7, 20, 29^. Even though the formulations of the individual ASM are distinct, experimental results from different formulations claim to provide physiologically relevant depictions of pathogen growth and physiology similar to that in CF sputum.^3, 7, 30^

Herein, we perform comparative metabolomics of the *P. aeruginosa* specialized metabolome grown in nine different ASM formulations. *P. aeruginosa* is the primary bacterial opportunistic pathogen infecting CF lungs and contributes to lung function decline.^31, 32^ In the CF lung, *P. aeruginosa* grows as biofilms, exopolysaccharide encased communities, which are recalcitrant to innate immune response and antibiotics.^33^ Within these biofilms, *P. aeruginosa* produces a suite of small molecule virulence factors that enable it to establish chronic infections by damaging host cells, outcompeting other members of the pulmonary microbiome, and developing as biofilms.^30, 34–36^ As *P. aeruginosa* specialized metabolites are associated with virulence phenotypes, their measurement can be used to understand how different nutrient availability in ASM formulations affect *P. aeruginosa* virulence.^37, 38^ We show that different ASM formulations induce differential production of *P. aeruginosa* PAO1 small molecule virulence factors, including members of the phenazine, quinolone, rhamnolipid, pyochelin, and pyoverdine molecular families, which is partially driven by the inclusion of commercial porcine gastric mucin (PGM) in the media.

## Results and Discussion

### The nutrient availability of different ASM formulations influences *Pseudomonas aeruginosa* PAO1 phenotype and secondary metabolic profile

To test the hypothesis that different ASM formulations would result in differential production of *P. aeruginosa* specialized metabolites, strain PAO1 was statically cultured in nine different ASM formulations.^3–8, 10, 11^ We chose the nine ASM formulations based upon citation count, prior use in our research, and if the formulation was a predecessor to other formulations. Broadly, these ASM can be separated into two lineages, complex and chemically defined. The complex lineage originates with Soothill ASM^4^, which was modified to yield Romling^9, 10^, Winstanley^6^, ASMDM^3^, Cordwell^5^, and SDSU^8^ ASM formulations (Figure 1). Soothill ASM was created using literature values for key nutrients identified in CF sputum, taking into account economic cost of materials.^4^ This base medium was modified by other researchers by addition of amino acids^6, 9, 10^, bovine serum albumin (BSA)^6^, alterations of mucin^3, 8^ and DNA^3^ concentration, and replacement of amino acid^5, 8^ or iron source.^5^ Concurrent to the development of the complex ASM lineage, the chemically defined synthetic CF medium (SCFM) series of ASM was created from the average concentrations of ions, free amino acids, glucose, and lactate measured from CF sputum samples (Figure 1).^7^ To more closely resemble the *in vivo* CF lung environment, the SCFM1 base medium was complemented with DNA, mucin, 1,2-Dioleoyl-sn-glycero-3-phosphocholine (DOPC) as a lipid source, and *N*-acetyl glucosamine (GlcNAc) as well as various nucleosides to create SCFM2 and SCFM3, respectively.^11^

**Figure 1.**
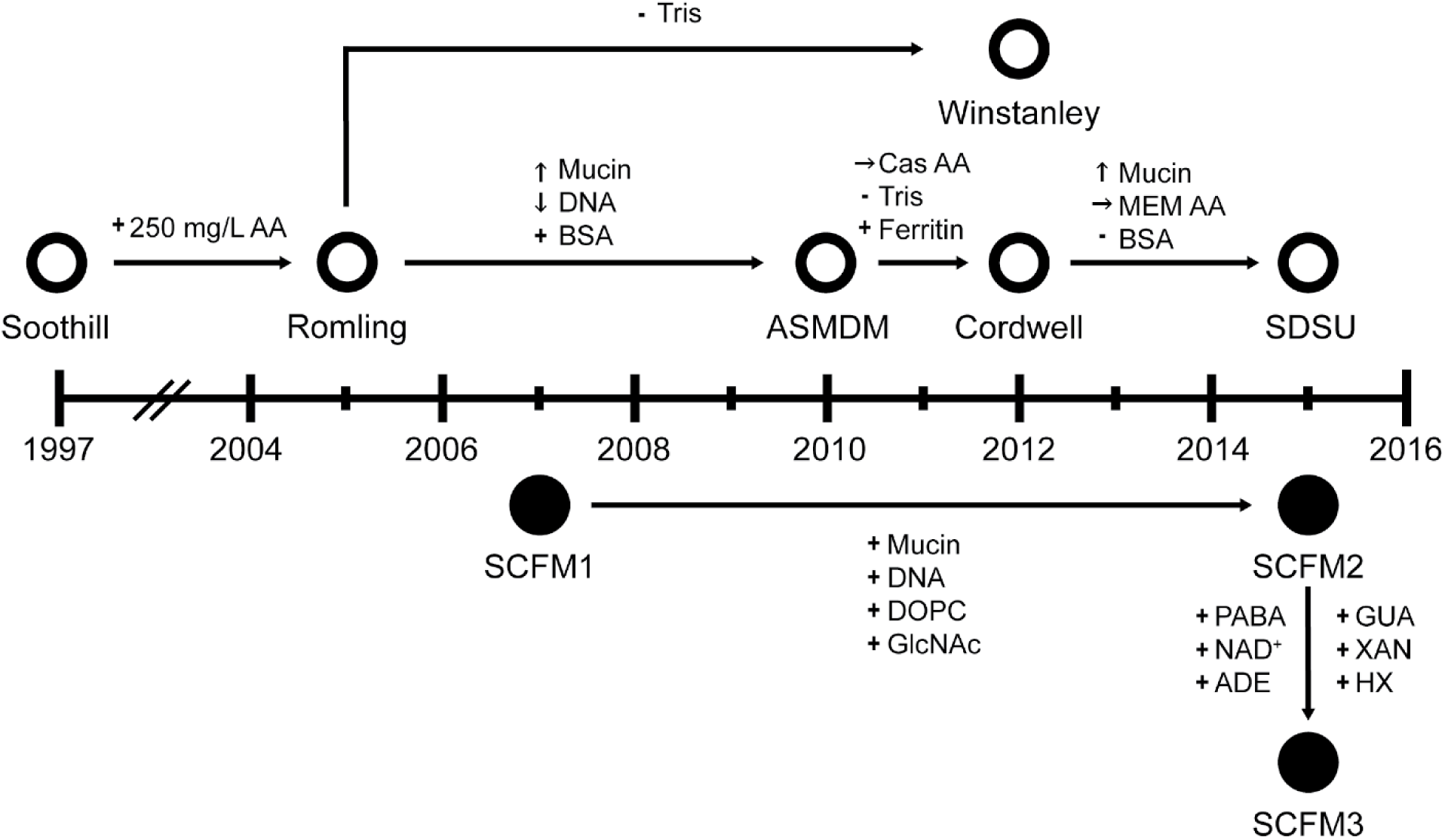
Publication timeline of ASM formulations. Each formulation is plotted at the year it was first published. Open circles indicate media derived from the Soothill ASM recipe. Closed circles indicate SCFM ASM lineage. Nutrient alterations between formulations are indicated by arrows. Compared to preceding formulation: + = added, - = removed, ↑ = increased, ↓ = decreased, → = different source. AA = amino acids, BSA = bovine serum albumin, Cas AA = casamino acids, MEM AA = MEM amino acids, DOPC = 1,2-dioleoyl-sn-glycero-3-phosphocholine, GlcNAc = *N*-acetyl glucosamine, PABA = 4-aminobenzoic acid, NAD+ = nicotinamide adenine dinucleotide, ADE = adenine, GUA = guanine, XAN = xanthine, HX = hypoxanthine. Recipes for ASM formulations are in detailed in Dataset S1 ASM formulations.

The gross morphology of the PAO1 biofilms and color of the cultures varied between media (Figure 2A), despite similar levels of growth (Figure 2B). The Soothill ASM PAO1 cultures were turbid, while the other eight ASM formulations induced the development of biofilm macrostructures. Addition of amino acids to the Soothill ASM to create Romling and Winstanley ASM led to the formation of a brown macrostructure phenotype by PAO1. Nutritional alterations of Romling ASM to create ASMDM, Cordwell, and SDSU ASM formulations led to assorted phenotypes of PAO1 growth, with structurally varied biofilms with diverse coloration. In ASMDM, *P. aeruginosa* grew heterogeneously with a layer of growth at the air-liquid interface, giving way to turbid growth further down the culture. In SDSU ASM, PAO1 formed a green, floating structure near the center of the well that reached the surface of the medium. PAO1 cultures in SCFM2 and SCFM3 were highly similar, with larger biofilm structures than in SCFM1 and a distinctly blue coloration, likely from high levels of pyocyanin.^39^

**Figure 2.**
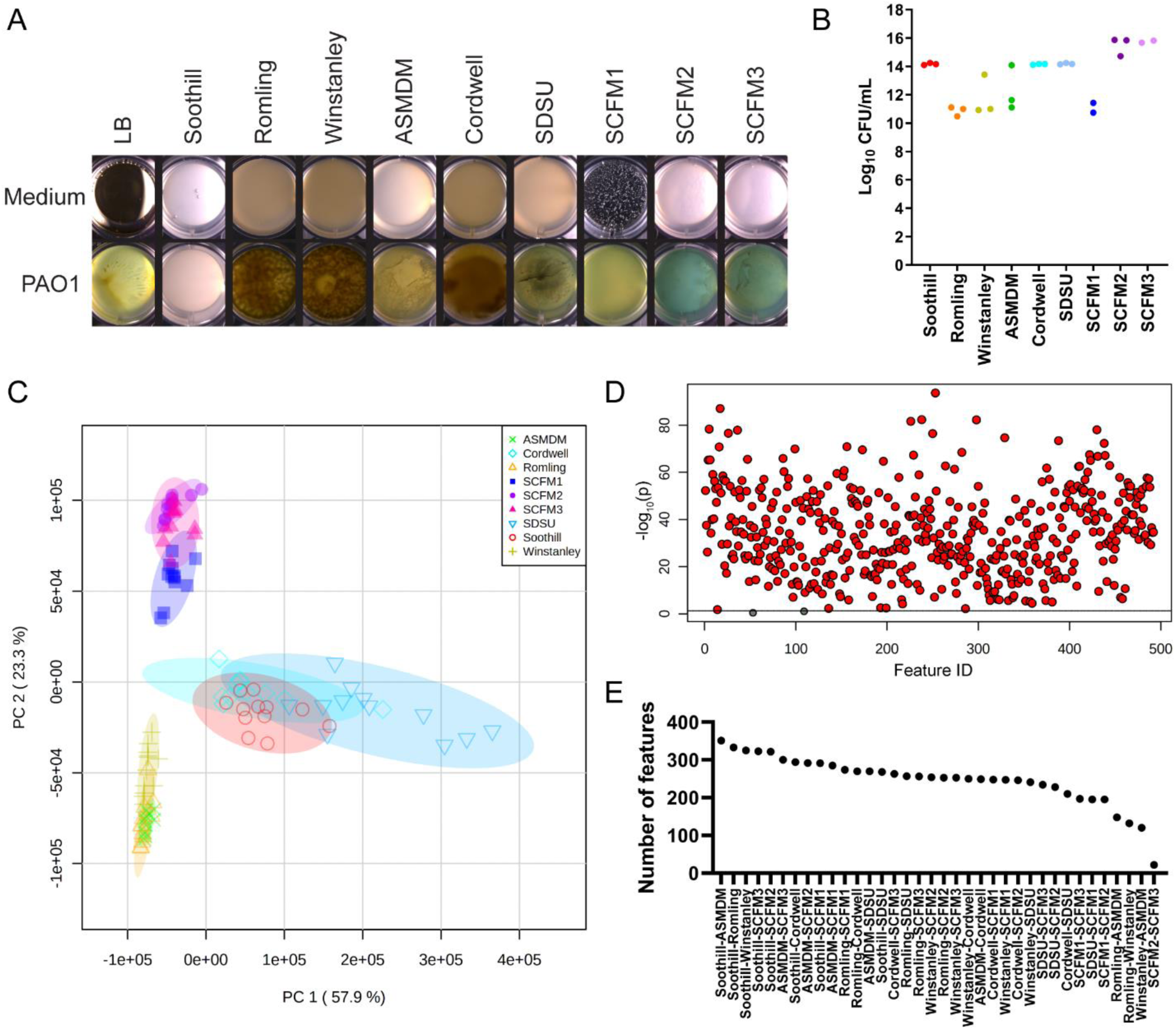
*P. aeruginosa* growth and global metabolism in ASM. (A) Phenotypes, (B) growth of representative replicates, (C) principal component analysis (PCA) score plots of metabolic profiles, (D) One-way ANOVA plot of metabolic features, and (E) pairwise number of statistically different metabolic features of PAO1 cultures in ASM formulations. Samples as labeled and colored as indicated. n=2-3 for CFU/mL growth assessment. n=12 for metabolite profiling. Statistically significant metabolic features (*p* < 0.05) are colored red in the one-way ANOVA plot.

As the color of *P. aeruginosa* cultures is largely dependent upon the abundance of its small molecular virulence factors, we applied comparative metabolomics to PAO1 cultures in the nine ASM formulations. Principal component analysis was used to visualize the similarity within and between the metabolome of PAO1 cultures in different ASM formulations (Figure 2C). Broadly, the metabolomes of biological replicates clustered together, with SDSU ASM cultures showing the highest between-replicate variability, likely due to the high concentration of mucin in the medium, which may affect metabolome extraction efficiency due to the high viscosity of the samples.^8^ In the PCA, samples clustered into three groups, largely based upon their formulations. The nutritionally distinct SCFM ASM formulations clustered separately from the Soothill ASM derived media. Within the Soothill ASM lineage samples, two distinct clusters were formed; one cluster encompassing Romling, Winstanley, and ASMDM ASM and the other containing Cordwell, SDSU, and Soothill ASM. The latter three ASM formulations have different amino acid content than the former (Figure 1), which likely affects the production levels of *P. aeruginosa* small molecule virulence factors, as amino acids are biosynthetic precursors of these metabolites^40–42^. Out of 490 molecular features detected, all but two were statistically significantly different between the samples (Figure 2D). The number of statistically different features produced in each pairwise interaction (Figure 2E) corresponds well to the PCA results, with cultures from nutritionally related media yielding the most similar metabolomes.

### ASM formulation variation stimulates differential production of small molecule virulence factors by PAO1

Since statistical analysis of the metabolomics data was unable to identify specific metabolic features that differentiated *P. aeruginosa* metabolism in the different ASM formulations, we used feature based molecular networking (FBMN) to annotate, quantify, and visualize the production of small molecule virulence factors produced by PAO1 in all nine ASM formulations.^43, 44^ *P. aeruginosa* has a well-characterized specialized metabolome, including phenazines, quinolones, rhamnolipids, and the siderophores pyoverdine and pyochelin.^37, 45, 46^ *P. aeruginosa* specialized metabolites have been detected in CF samples^30, 34, 35, 47–53^ and are associated with pathogenicity and virulence.^29, 37, 38^ As nutrient concentration varies between ASM formulations, alteration in production levels of these metabolites by *P. aeruginosa* may influence interpretation of virulence-related phenotypes.

Phenazines are redox-active metabolites with various biological activities, including roles in promoting iron uptake^54, 55^, anaerobic respiration^54^, biofilm formation^56, 57^, and interspecies interactions.^45, 55, 58, 59^ We quantified the production of four phenazines: 1-hydroxphenazine (1-HP), pyocyanin (PYO), phenazine-1-carboxamide (PCN), and phenazine-1-carboxylic acid (PCA) (Figure 3). Two metabolic features have matches to the GNPS spectral libraries for PYO (Figure 3A). These features have different chromatographic retention times, but identical MS/MS fragmentation patterns, suggesting they are structurally related. While 1-HP, PCA, and PYO have been detected in CF sputum^30, 52^, their detection across samples has been spurious and UV detection may not be sufficient to differentiate phenazine production from other components of sputum.^60^ Therefore, phenazine production is most commonly measured from pure cultures of CF clinical isolates.^61^ Phenazines were quantified from all PAO1 cultures. However, the abundance of specific phenazines (Figure 3A), total phenazine production (Figure 3B), and the composition of the phenazine molecular family varied between ASM formulations (Figure 3C). Phenazine production was lowest in Soothill ASM and highest in SCFM2 and SCFM3 (Figure 3B). Soothill ASM does not contain amino acids,^4^ restricting the amino acid precursor availability for the biosynthesis of siderophores by *P. aeruginosa*.^62–64^ In this medium, PAO1 may produce a higher proportion of PCA relative to other phenazines (Figure 3D) in order to acquire iron from the media as PCA has been shown to reduce Fe^3+^ to Fe^2+^ in a *P. aeruginosa* mutant unable to produce pyochelin and pyoverdine.^65^ PYO is produced at the highest levels by PAO1 grown in SCFM2 and SCFM3 (Figure 3A/C), which contributes to the blue phenotype of these cultures (Figure 2A).^39^ SCFM2 and SCFM3 are compositionally different from SCFM1 by the addition of several components, including *N-*acetyl glucosamine (GlcNAc).^11^ Addition of GlcNAc to SCFM1 was shown to increase production of PYO by *P. aeruginosa*, likely through a mechanism to sense and respond to the presence of peptidoglycan.^66^

**Figure 3.**
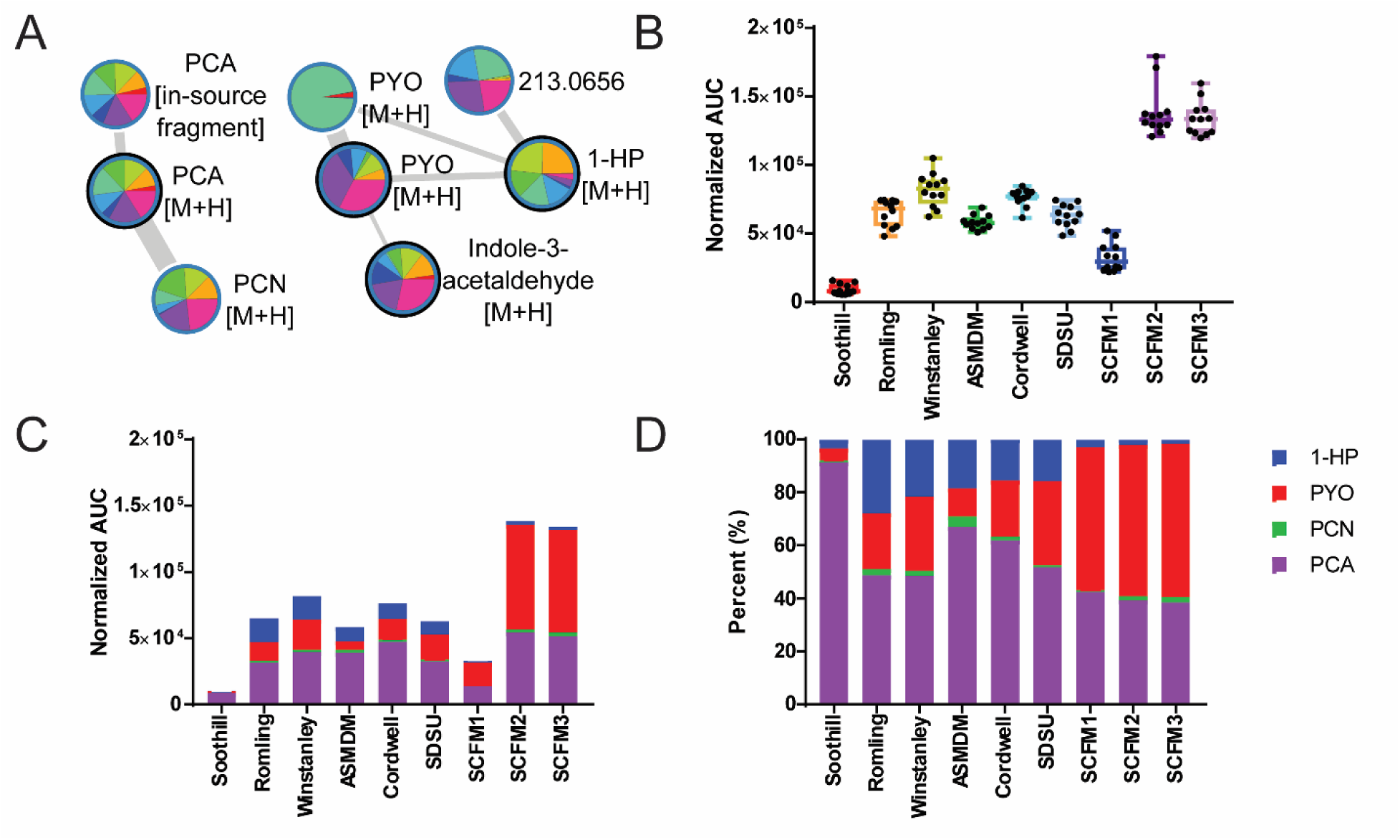
Phenazine production by PAO1 in ASM. (A) Phenazine molecular family visualized with feature based molecular networking, colored as in (B). (B) Total normalized area under the curve (AUC) for the phenazine molecular family in each ASM formulation. Box plots represent the 25-75^th^ percentile, with a line at the median. Error bars indicate the minimum to maximum. Individual samples values shown (n=12 per ASM formulation). (C) Profile of total phenazine production, colored as in (D). (D) Proportion of total phenazines. 1-HP: 1-hydroxyphenazine; PYO: pyocyanin; PCN: phenazine-1-carboxamide; PCA: phenazine-1-carboxylic acid.

*P. aeruginosa* is known to produce over 50 quinolones^67^ in three structural sub-classes based on the characterized molecules: 2-heptyl-4-quinolone (HHQ)^68^, 2-heptyl-3,4-dihydroxyquinoline (Pseudomonas Quorum Signal, PQS)^69^ and 2-heptyl-4-hydroxyquinoline N-oxide (HQNO).^70^ Different quinolones have different roles in intra- and interspecies interactions within the CF lung environment.^71–75^ HHQ and PQS are important quorum sensing signaling metabolites participating in regulating the production of a large suite of virulence factors, including PYO, and many genes related to iron limitation.^68, 69, 76–78^ HQNO lacks regulatory activity, but is an important metabolite in inter-species interactions with *Staphylococcus aureus* through inhibition of the respiratory chain^71, 72^ and enhancement of antibiotic tolerance.^79–81^ While we quantified 43 quinolones from the PAO1 cultures in ASM (Figure 4A), only 13 account for 85-95% of the total quinolones produced in ASM. Like phenazine production, levels of individual and total quinolones produced by PAO1 varied between ASM formulations, with PAO1 cultured in Romling, Winstanley, and ASMDM ASM formulations producing the highest levels of quinolones (Figure 4B). These three formulations have the highest concentration of the aromatic amino acids tryptophan, tyrosine, and phenylalanine. Importantly, tryptophan levels are low in CF sputum;^7^ therefore, these media may not replicate the amino acid availability *in vivo.* Higher concentrations of tryptophan may alter quinolone production as it is catabolized via the kynurenine pathway to generate anthranilate, a precursor of quinolone biosynthesis.^42, 82^ Both tyrosine and phenylalanine are important carbon sources in CF sputum and have been shown to induce PQS biosynthesis in SCFM1.^7, 83^

**Figure 4.**
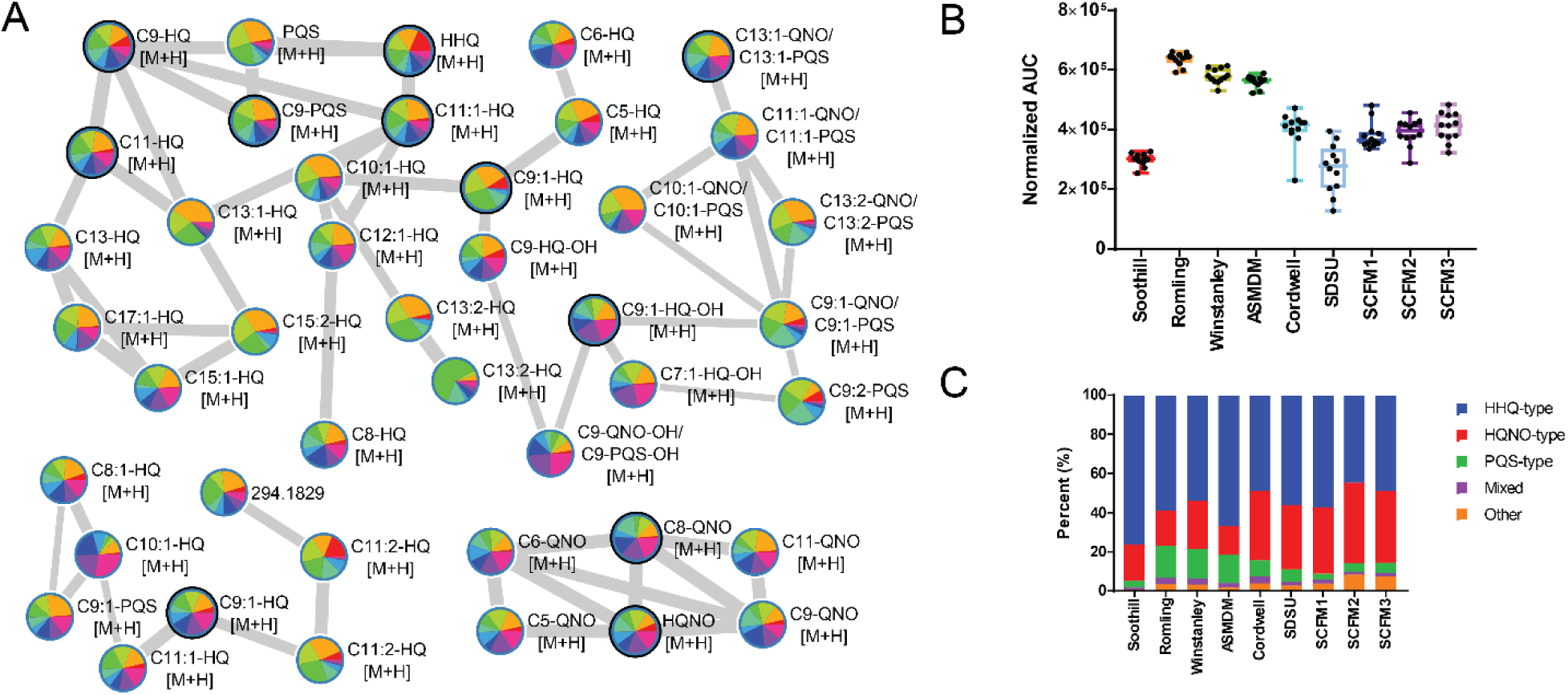
Quinolone production by PAO1 in ASM. (A) Quinolone molecular family visualized with feature based molecular networking, colored as indicated in (B). Nodes in the network are annotated by alkyl chain length and unsaturation, followed by structural sub-class assignment. HQ, PQS, and QNO denotes structural relatedness to HHQ, PQS, and HQNO, respectively. Features representing metabolites that could not be distinguished based upon MS/MS fragmentation patterns are indicated and referred to as ‘mixed’ in (C). Annotations with –OH designation indicate hydroxylation at unknown position within the molecule and are referred to as ‘other’ in (C). (B) Total normalized area under the curve (AUC) for the quinolone molecular family in each ASM formulation. Box plots represent the 25-75^th^ percentile, with a line at the median. Error bars indicate the minimum to maximum. Individual samples values shown (n=12 per ASM formulation). (C) Proportion of total quinolones based upon structural sub-class. Quinolone structural sub-classes are colored as indicated. HHQ: 2-heptyl-4-quinolone; PQS: 2-heptyl-3,4-dihydroxyquinoline; HQNO: 2-heptyl-4-hydroxyquinoline *N*-oxide.

In all ASM, PAO1 produced predominantly HHQ-type quinolones (44-76%), while HQNO-type were produced at 15-41% total quinolones, and PQS-type were the consistently the lowest abundant quinolones (3-17%) (Fig. 4C). The most abundant alkyl chain lengths were heptyl (C7), nonyl (C9), and nonenyl (C9:1), accounting for 33-60%, 24-41%, and 10-21% of the quinolones produced by PAO1 in each ASM, respectively. HHQ- and HQNO-type quinolones have been suggested for use as diagnostic biomarkers of *P. aeruginosa* infection in adults and children with CF^50^ as they are the most abundant quinolone congeners in sputum^30, 50, 84, 85^, explant lung tissue^86, 87^, BALF^50^, and serum^50^. Importantly, ASM formulation affects the ratio of C7, C9:1, and C9 quinolones produced by PAO1, most prominently for HHQ- and HQNO-type quinolones. For example, PAO1 cultured in Soothill ASM produces 2.5-fold more C7-HQ (HHQ) than C9-HQ (NHQ), while in SCFM2 PAO1 produces similar levels of both HHQ-type quinolones. It is unknown whether the C9 and C9:1 quinolone congeners have different biological function than their C7 analogues. The difference in quinolone production specificity between Soothill and SCFM ASM lineages may be the result of different sources of lipid precursors. Soothill-derived ASM includes egg yolk emulsion as a lipid source, while SCFM-derived ASM include a single lipid species (1,2-Dioleoyl-sn-glycero-3-phosphocholine, DOPC), although commercial PGM contains lipids as well (Figure 8).

Rhamnolipids are important virulence-associated glycolipids with functional roles in biofilm formation and maintenance^88, 89^, evasion of host immune response^90, 91^, competition with fungal neighbors^92^, transport of PQS^93^, and as biosurfactants^94^. Rhamnolipids have been detected in CF sputum^30, 47, 84, 85^ as well as explant lung tissue^86, 87^, and rhamnolipid production levels are higher in some virulent *P. aeruginosa* clinical isolates.^38^ Structurally, *P. aeruginosa* rhamnolipids consist of 1-2 rhamnose units bound to two fatty acids, yielding mono- and di-rhamnolipids, respectively. We quantified six rhamnolipids, three mono-rhamnolipids and three di-rhamnolipids (Figure 5A). Unlike phenazine and quinolone production by PAO1 in the different ASM formulations, rhamnolipid production was somewhat consistent across most cultures, with lowest levels of production by PAO1 in SCFM2 and SCFM3 (Figure 5B). The proportion of mono-to di-rhamnolipids produced by PAO1 was maintained across all ASM cultures (Figure 5C), with di-rhamnolipids accounting for 56-70% of total rhamnolipids. The predominant fatty acid chain length was C10, accounting for 56-70% of the rhamnolipids produced by PAO1 in ASM, which reflects the predominant rhamnolipids detected from CF sputum^30, 84, 85^ and explant lung samples.^86, 87^

**Figure 5.**
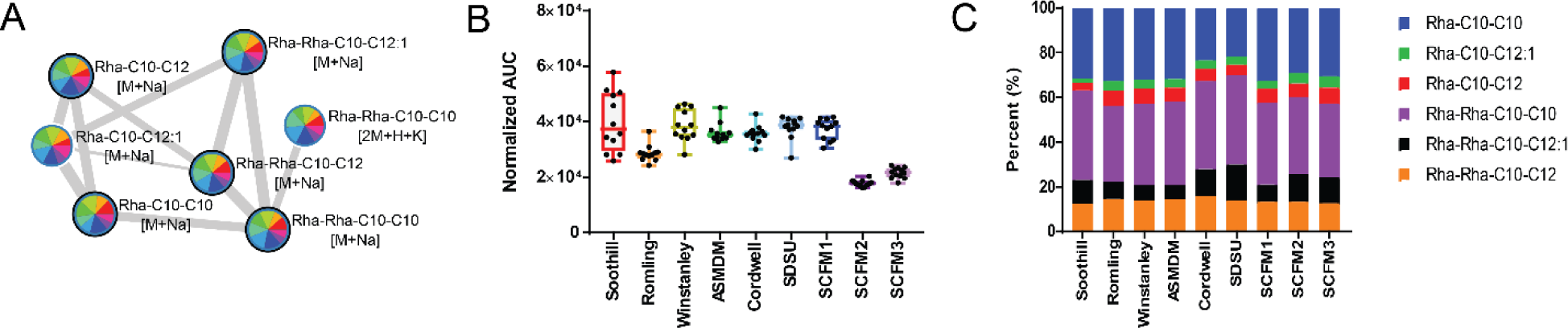
Rhamnolipid production by PAO1 in ASM. (A) Rhamnolipid molecular family (sodiated) visualized with feature based molecular networking, colored as indicated in (B). Nodes in the network are annotated by rhamnose units (Rha) followed by fatty acid chain lengths. Only sodium adducts are shown. (B) Total normalized area under the curve (AUC) for the quinolone molecular family in each ASM formulation. Box plots represent the 25-75^th^ percentile, with a line at the median. Error bars indicate the minimum to maximum. Individual samples values shown (n=12 per ASM formulation). (C) Proportion of total rhamnolipid production, colored as indicated.

The most striking difference in specialized metabolite production by PAO1 in different ASM formulations was the production of the siderophores pyochelin and pyoverdine, which were produced by PAO1 in SCFM1 cultures (Figure 6). While pyochelin was annotated by a MS/MS spectral match to the GNPS libraries^43^, the nodes corresponding to the pyoverdine molecular family were prioritized for annotation based upon coloration of the pie chart, indicating high production by PAO1 in SCFM2 (Figure 6A). Siderophores are metal scavenging virulence factors^95, 96^ secreted in low iron conditions by microbes to acquire environmental iron.^97–100^ Pyoverdine is considered the primary siderophore of *P. aeruginosa* as it has a higher affinity for iron than pyochelin, displaces iron from host proteins, and is required for virulence in acute infection models.^95, 96, 98, 101–103^ However, pyochelin production by *P. aeruginosa* typically occurs first as pyoverdine production is only initiated by *P. aeruginosa* when iron concentrations are extremely low.^104^ Both pyochelin and pyoverdine have been detected in CF sputum^84, 85^, with production of pyoverdine highly dependent on strain^34, 105^. Although pyoverdine biosynthesis was reported to be upregulated in SCFM1 by *P. aeruginosa*^7^, we were surprised to detect pyoverdines as the production of this molecular family by PAO1 is below our limit of detection when cultured in most media. The high abundance of pyochelin and pyoverdine from SCFM1 PAO1 cultures suggests that only this ASM formulation is iron limited, which does not align with the published media compositions. Prior to addition of the iron chelator diethylenetriaminepentaacetic acid (DPTA) to the media, Soothill ASM was measured to have an Fe^2+^ concentration of 9.5 µM^4^, and the same iron concentration was presumed for Romling, Winstanley, and ASMDM ASM formulations. Both Cordwell and SDSU ASM contain 0.000003% (w/v) ferritin^5, 8^, which adds ∼2.85 µM Fe^3+^ to the media. Last, the SCFM series of ASM all include 3.60 µM Fe^2+^ from addition of iron sulfate.^7, 11^ As SCFM2 nutrient availability differs from SCFM1 by only the addition of GlcNAc, DNA, DOPC, and mucin^11^, these results suggest that one of these additives is an undefined source of iron.

**Figure 6.**
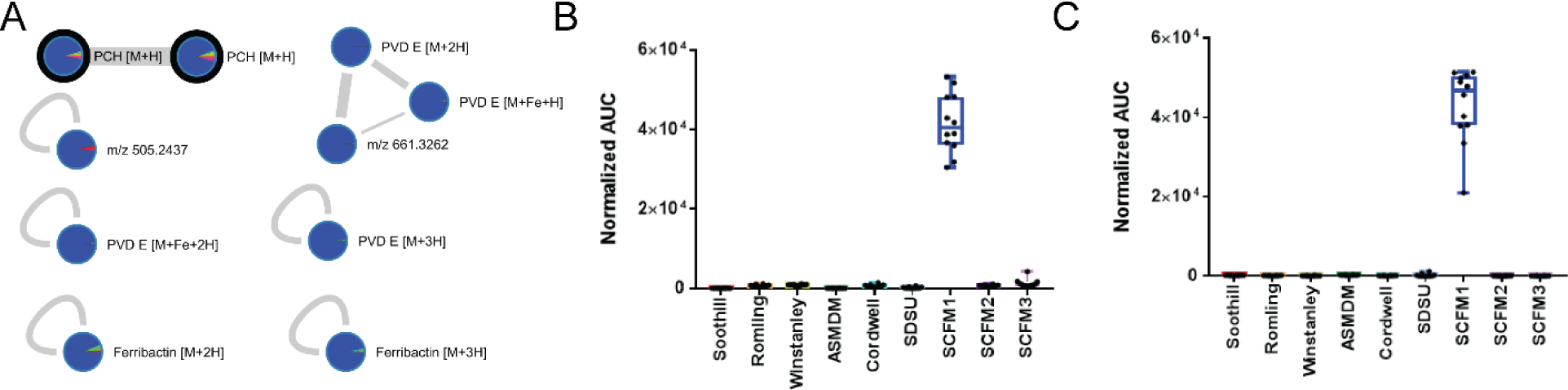
Siderophore production by PAO1 in ASM. (A) Pyochelin and pyoverdine molecular families visualized with feature based molecular networking. SCFM1 cultures are blue. Nodes in the network are annotated by adducts. (B) Total normalized area under the curve (AUC) for pyochelin in each ASM formulation (n=12 per ASM formulation). (C) Total normalized AUC for pyoverdine molecular family in each ASM formulation (n=12 per ASM formulation). For (C) and (D), box plots represent the 25-75th percentile, with a line at the median. Error bars indicate the minimum to maximum.

### Commercial porcine gastric mucin (PGM) contains contaminants which affect PAO1 specialized metabolite production

As mucin was the most complex additive to our SCFM2 media, we hypothesized that it was a potential source of contaminating iron. Mucins are high molecular weight glycoproteins responsible for the viscoelastic properties of mucus.^106, 107^ In CF, loss of function of the cystic fibrosis transmembrane conductance regulator (CFTR) leads to altered ion transport and accumulation of thick bronchial mucus in the lungs.^25, 108^ The altered biophysical properties of pulmonary mucus in CF contributes to the establishment of chronic infections by pathogens, including *P. aeruginosa*.^109–111^ Due to its importance in CF pathogenesis, inclusion of mucin in ASM is critical to fully replicating the *in vivo* nutritional^26, 112, 113^ and structural^112, 114–116^ CF pulmonary environment.^13, 116, 117^ However, despite the limitations associated with altered physical and chemical properties of the mucins associated with the purification process and inclusion of a number of mucin-bound impurities^118–120^, the most economical choice is commercial porcine gastric mucin (PGM), which is often used as a component of ASM without further purification.

Indeed, SCFM1 complemented with 0.5% (w/v) PGM (PGM-SCFM1) was sufficient to alter production of the phenazine, quinolone, and rhamnolipid molecular families as well as the siderophores, pyochelin and pyoverdine by PAO1 compared to SCFM1 (Figure 7). Overall phenazine production was increased in PAO1 cultures in PGM-SCFM1 (Figure 7A), with increased abundance of 1-HP, PYO, and PCA. Unlike phenazine abundance, quinolone production by PAO1 was similar in the two media (Figure 7B). However, the composition of the quinolone molecular family was altered with PGM addition, with an increase in C7-HQ (HHQ) and a decrease in C9:1-HQ-OH and C9-QNO (NQNO). Importantly, PQS-type quinolones were not detected in this experiment. Surprisingly, although HHQ production increased with PGM addition, C9-HQ (NHQ) production did not. Conversely, while C9-QNO (NQNO) production decreased, C7-QNO (HQNO) production remained the same. Rhamnolipid production was decreased in PGM-SCFM1 PAO1 cultures (Figure 7C). Lastly, addition of PGM to SCFM1 was sufficient to reduce siderophore production by PAO1 (Figure 7D). The changes in specialized metabolite production by PAO1 in PGM-SCFM1 compared to the SCFM1 cultures parallels the differential production of these metabolites between SCFM1 and SCFM2 cultures, indicating that the other additives are minimally contributing to these alterations.

**Figure 7.**
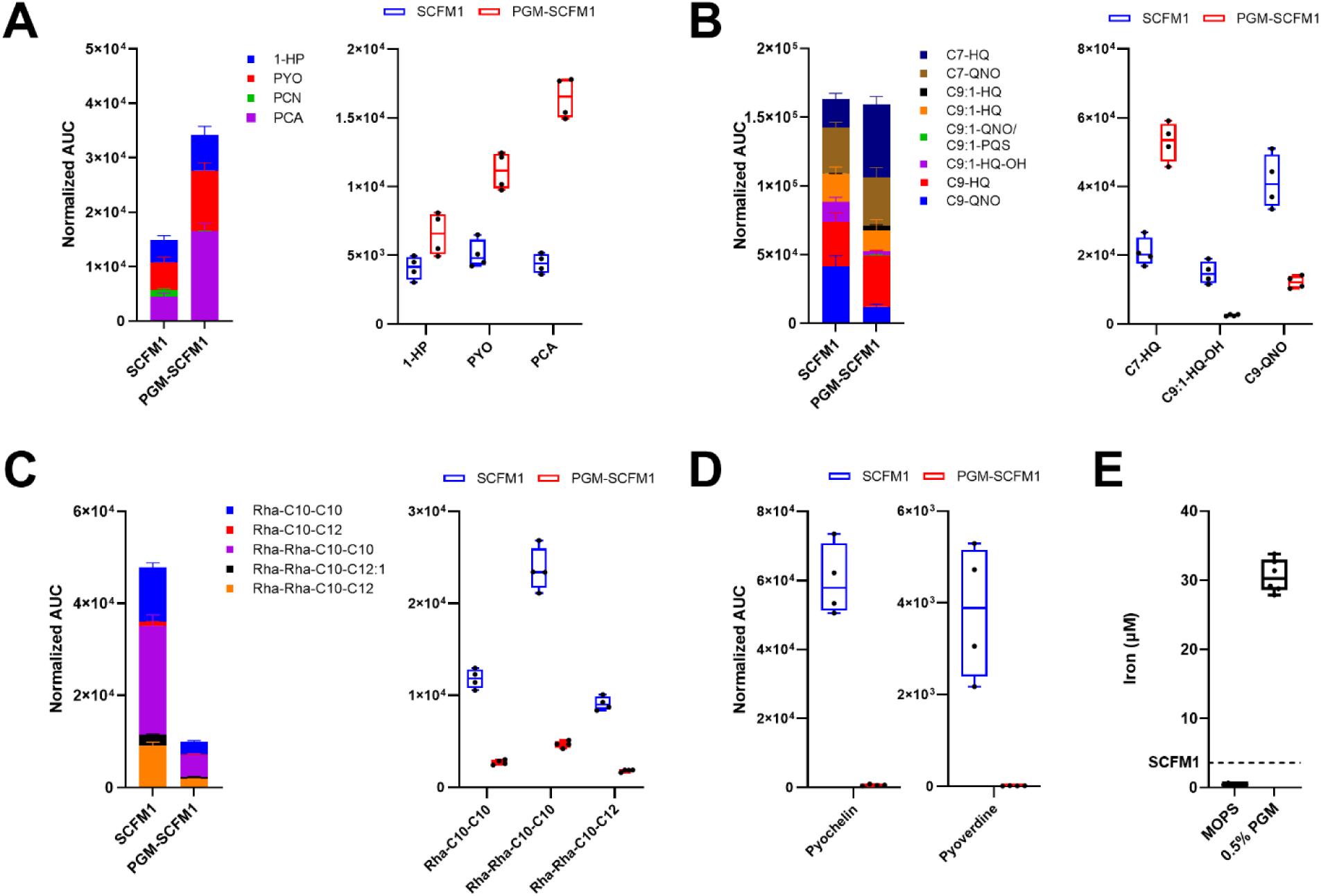
Effect of 0.5% PGM on production of small molecule virulence factors by PAO1. (A) Left: Total normalized AUC for phenazine molecular family, colored as indicated. Right: Normalized AUC for phenazine molecular family members. (B) Left: Total normalized AUC for phenazine molecular family, colored as indicated and only most abundant quinolones included. Right: Normalized AUC for quinolone molecular family members with largest alterations in production. (C) Left: Total normalized AUC for rhamnolipid molecular family, colored as indicated. Right: Normalized AUC for rhamnolipid molecular family members with largest alterations in production. (D) Normalized AUC for pyochelin and pyoverdine molecular family. (E) Concentration of iron in 0.5% PGM and vehicle control (MOPS). Dashed line indicates the concentration of iron included in the SCFM1 ASM formulation. For (A)-(D), n=4 biological replicates. For all stacked bar charts, bars represent the mean and error bars indicate standard deviation of the mean. All box plots represent the 25-75th percentile, with a line at the median, and error bars indicating the minimum and maximum.

As the composition of commercial mucins is largely undefined, the alterations in production of small molecule virulence factors by PAO1 cultured in PGM-SCFM1 could be due to the presence of mucin, assorted iron sources, GlcNAc, or another component in the PGM. Confirming our suspicion that commercial PGM contains iron, ICP-MS analysis revealed 0.5% PGM contains approximately 30 μM iron (Figure 7E). The abundance of all metals quantified from PGM are summarized in Figure 8A. The concentration of iron in commercial PGM is 8.5 times higher than the concentration of iron added to SCFM1 medium (3.6 μM). As ASMDM and Cordwell ASM formulations contain 1.0% (w/v) PGM and SDSU ASM contains 2.0% (w/v) PGM, the levels of iron in these media correspond to two and four times higher than the other ASM formulations, respectively. Inclusion of PGM in SCFM1 increases the concentration of iron in the medium to a more clinically relevant level of iron.^121–123^ However, the distinct iron sources in PGM are unknown and may not represent those in the CF lung.^122, 124^

**Figure 8.**
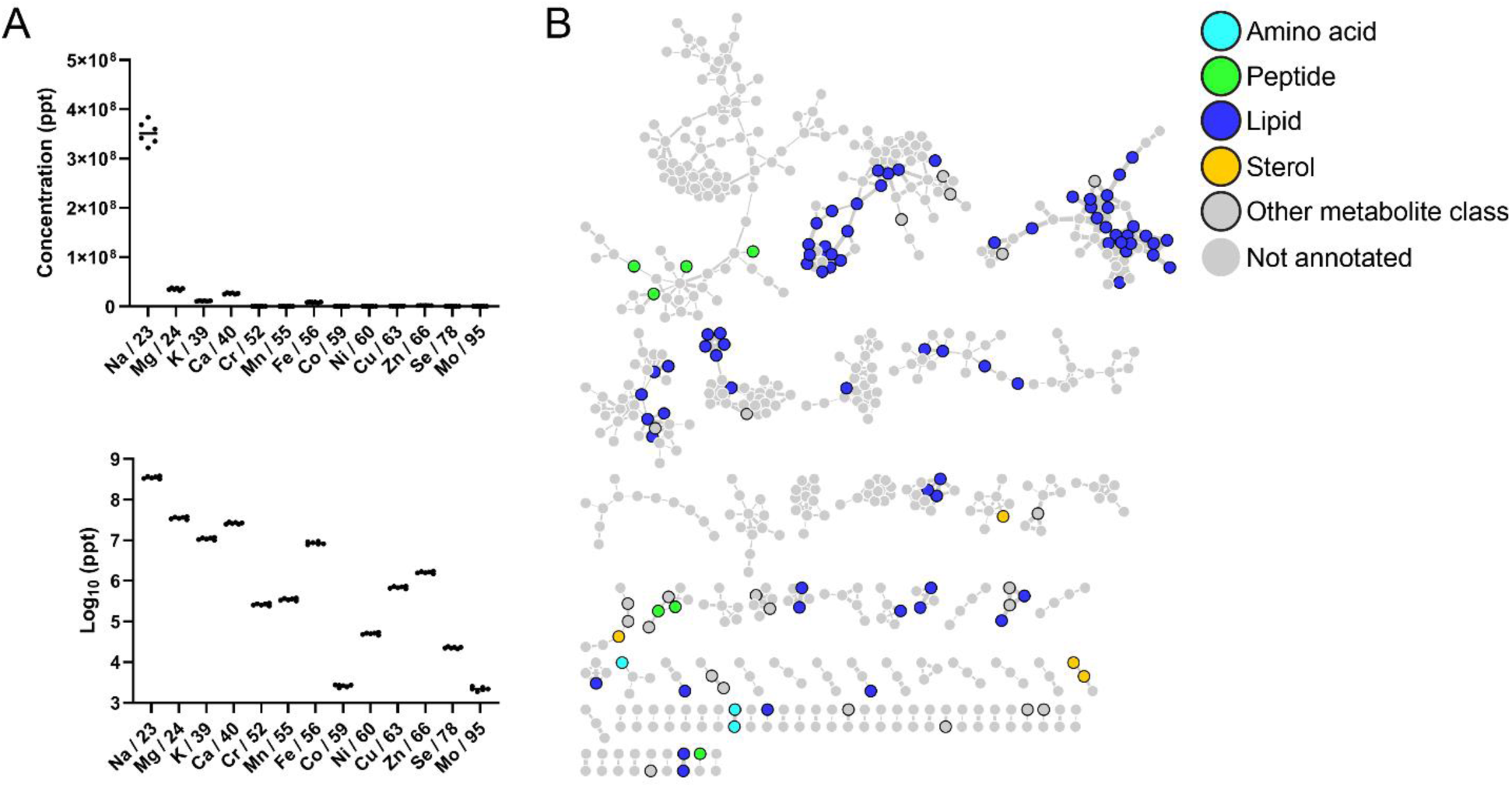
Contaminants in Commercial PGM. (A) Metals detected in PGM. Line indicates the median of six technical replicates. (B) Classical molecular network of lipophilic extracts of commercial PGM. Colored as indicated.

Due to the role of iron concentration in regulating specialized metabolite production by *P.* aeruginosa, we are confident that the high concentration of iron contributed by PGM is affecting the production levels of these metabolites in PGM-SCFM1.^89, 125, 126^ However, during analysis of the FBMN of SCFM1 and PGM-SCFM1 cultures, several molecular features matched to lipid spectra in the GNPS libraries, which could only be present in the PGM, as they were not separately added to the media. Analysis of lipophilic extracts of commercial PGM using classical molecular networking (CMN) revealed that in addition to contamination with a variety of metals, PGM contains amino acids, small peptides, and lipids (Figure 8B). Although SCFM1 is tryptophan limited, tryptophan was detected in PGM. Therefore, addition of PGM to the medium increases the tryptophan concentration, which can serve as a source of anthranilate in quinolone biosynthesis.^127^ Like amino acids, bioactive host lipids may play important roles in shaping CF lung ecology, including eicosanoids, sphingolipids, and ceramides.^128, 129^ PGM contains a wide variety of lipids (Figure 8B), including those most highly detected in CF sputum samples.^30, 128^ Detection of these contaminants within commercial PGM suggest that inclusion of crude PGM in culture medium will likely yield null results when investigating the mechanistic roles of these metabolites (as well as those not detected under our extraction conditions) on microbial physiology, virulence, and ecology.

Further research will be required to establish what specific components of commercial PGM are responsible for alteration in PAO1 specialized metabolite production and to develop cost-effective methods for purification of mucin from crude preparations. The mucin concentration of CF sputum ranges from 16-30 mg/mL.^25^ While methods have been established for isolating pure PGM from pig stomach, only ∼65 mg of PGM can be purified from a single stomach.^118^ To produce 1 L of an ASM formulation containing 0.5% (w/v) mucin (5 mg/mL), this purification would require processing of at least 77 pig stomachs. Other laboratories have established procedures for purifying crude commercial mucins^130^, but without performing analytical analysis to measure all potential contaminants, it is impossible to assess the purity of the mucins.

## CONCLUSION

*In vitro* model systems are attractive due to their broad accessibility and ability to probe hypotheses thoroughly. The foundation of any *in vitro* investigation is the medium. In CF, a broad suite of different ASM formulations has been created. In this study, we demonstrated that differences in nutrient content between ASM formulations alter the phenotype and metabolomics profile of *P. aeruginosa* PAO1. Importantly, the concentration of specialized metabolites, including members of the phenazine, quinolone, rhamnolipids, and siderophore molecular families, are impacted by ASM formulation. Given the role of these metabolites in virulence, differences in ASM formulation may influence the overall outcome of interactions of *P. aeruginosa* with host cells and other members of the CF pulmonary microbiota. Commercial PGM was identified as a source of contaminating bioavailable metals, lipids, amino acids, and small peptides. Addition of commercial PGM to the chemically defined SCFM1 ASM formulation was sufficient to alter production of small molecule virulence factors by PAO1. In view of the importance of mucin in replicating mucosal infection environments *in vitro,* further work is needed to understand the contribution of individual components of commercial mucin to microbial physiology and virulence, as well as development of cost-effective means to purify mucins from crude commercial preparations.

## MATERIALS AND METHODS

### General

*Pseudomonas aeruginosa* PAO1 (MPAO1) was acquired from the University of Washington, Seattle^113, 131^. All media components, concentrations, and commercial sources for artificial sputum media (ASM) formulations are listed in **Dataset S1 ASM formulations**^3–11^. Miller Luria broth (LB) and porcine gastric mucin type III (PGM) were purchased from Millipore Sigma. Organic solvents for extraction and liquid chromatography tandem mass spectrometry (LC-MS/MS) analysis were purchased from VWR (ethyl acetate (EtOAc), HiPerSolv Chromanorm purity) and Fisher Scientific (methanol (MeOH), acetonitrile (ACN), formic acid (FA), and water; all Optima LC/MS grade). PA Mix was created as a set of external standards for *P. aeruginosa* specialized metabolite annotation. PA Mix contains 10 µM of each of the following: 1-hydroxyphenazine (1-HP, Tokyo Chemical Industry); pyocyanin (PYO, Sigma); phenazine-1-carboxylic acid (PCA, Ark Pharm); phenazine-1-carboxamide (PCN, Ark Pharm); 2-heptylquinolin-4(1H)-one (HHQ, Ark Pharm); 2-heptyl-4-hydroxyquinoline *N*-oxide (HQNO, Cayman Chemical); 2-heptyl-3-hydroxyl-4-quinolone (Pseudomonas Quinolone Signal, PQS, Chemodex); N-butyryl-l-homoserine lactone (C4-HSL, Cayman Chemical); N-(3-oxododecanoyl)-l-homoserine lactone (3-oxo-C12-HSL, Millipore Sigma); and a rhamnolipid mixture (RHL, AGAE Technologies, 90% pure).

### Identification of commonly used ASM formulations

The Web of Science Core Collection (Clarivate Analytics) was searched for papers with the ‘TOPIC’ terms: “sputum medium”, “sputum media”, “cystic fibrosis medium”, “cystic fibrosis media”, “sputa media”, “sputa medium”, “CF media”, or “CF medium” in the title, abstract, or keywords fields on May 10, 2020. All terms were searched as shown, including quotation marks. The results of this search were analyzed using the Web of Science tools ‘Create Citation Report’ and ‘Analyze Results’. The Analyze Results tool was used to download data corresponding to Web of Science Categories. The citation report was used to select ASM formulations that were commonly cited.

### Preparation of media

LB broth was prepared according to manufacturer instructions. Commercial PGM was suspended in 1X 3-morpholinopropane-1-sulfonic acid (MOPS) buffer (pH 7.0) to make a 10% (w/v) suspension, sterilized using a liquid autoclave cycle (<20 minute sterilization time), and an aliquot was checked for sterility before use. ASM formulations were prepared as reported^3–11^ with the following modifications. Antibiotics were not included in any medium. For the Cordwell formulation^5^, the reported 0.1% (w/v) agar was excluded. For SCFM2 and SCFM3^11^, bovine submaxillary mucin (BSM) was replaced with PGM at the published concentration, a common substitution^12–14^. Each ASM formulation was brought to the published pH^3–11^. All media was stored in the dark at 4°C, checked for sterility prior to use, and used within one month of preparation.

### Cultivation of *P. aeruginosa* in LB and ASM formulations

PAO1 was inoculated from a streak plate into 5 mL LB broth and incubated overnight at 37°C, shaking at 220 RPM. 1.98 mL of LB or ASM were inoculated with 20 µL of *P. aeruginosa* LB broth culture (OD600 of 0.05; ∼10^6^ CFU/mL final concentration) in polystyrene 24 well plates^6^. For assessing *P. aeruginosa* growth and specialized metabolite production, each medium was evaluated in its own 24 well plate (12 medium control wells and 12 growth wells per medium). The media were inoculated and processed in three batches, comprised of 2-4 ASM formulations and a parallel LB plate as a control. All plates were covered and incubated statically at 37°C for 72 hours. For assessing the addition of PGM to SCFM1 on *P. aeruginosa* growth and specialized metabolite production, *P. aeruginosa* was cultured in SCFM1 and PGM-SCFM1 in a single polystyrene 24 well plate (4 medium control wells and 4 growth wells per medium).

### Processing of *P. aeruginosa* cultures in LB and ASM formulations

Gross phenotypes of *P. aeruginosa* growth were photographed using a top-down view with a Stemi 508 stereo microscope with an Axiocam 105 color camera (Zeiss). Following visualization, 100 µL cellulase (*Aspergillus niger*, 100 mg/mL in sterile water; Sigma) was added to each well and aggregates were mechanically disrupted by pipetting using wide-bore pipette tips (USA Scientific), followed by water bath sonication (Branson) for 10 minutes, and a secondary mechanical disruption by pipetting using wide-bore pipette tips. Disrupted samples were aliquoted for measurements of growth (CFUs) and metabolomics analysis. Three representative biological replicates from each culture condition were serially diluted, spotted onto LB agar, incubated at 37°C overnight, and counted to determine CFU/mL.

### Processing of *P. aeruginosa* cultures in SCFM1 and PGM-SCFM1

Aggregates were mechanically disrupted by pipetting using wide-bore pipette tips (USA Scientific). Disrupted samples were aliquoted for measurements of growth (CFUs) and metabolomics analysis. All four biological replicates from each culture condition were serially diluted, spotted onto LB agar, incubated at 37°C overnight, and counted to determine CFU/mL.

### ASM sample preparation for LC-MS/MS

Each sample aliquot was chemically disrupted with an equal volume of 1:1 EtOAc:MeOH. The samples were dried, resuspended in 100% MeOH containing 1 µM glycocholic acid (Calbiochem, 100.1% pure), diluted as needed in 100% MeOH containing 1 µM glycocholic acid, and centrifuged for 10 min at 4,000 RPM (Thermo Sorvall ST 40R) to remove non-soluble particulates. Samples were diluted as follows for measurement of *P. aeruginosa* specialized metabolite production in ASM formulations and LB: not diluted: Soothill ASM; 5-fold dilution: Romling ASM, Cordwell ASM, SCFM1, SCFM2, and SCFM3; 10-fold dilution: Winstanley ASM, ASMDM, SDSU ASM, and LB. All samples for measurement of *P. aeruginosa* specialized metabolite production in SCFM1 and PGM-SCFM1 were diluted 10-fold.

### PGM sample preparation for LC-MS/MS

50 µL of 10% (w/v) PGM in 1X MOPS was chemically extracted in technical triplicate with 200 μL of 1:1 EtOAc:MeOH and following the Bligh-Dyer extraction protocol.^132^ The samples were dried and resuspended in 100% MeOH containing 1 μM glycocholic acid, centrifuged for 10 min at 4,000 RPM, and diluted 10-fold.

### LC-MS/MS data collection

Mass spectrometry data acquisition for all samples was performed using a Bruker Daltonics Maxis II HD qTOF mass spectrometer equipped with a standard electrospray ionization (ESI) source. The mass spectrometer was tuned by infusion of Tuning Mix ESI-TOF (Agilent Technologies) at a 3 µL/min flow rate. For accurate mass measurements, a wick saturated with Hexakis(1H,1H,2H-difluoroethoxy) phosphazene ions (Apollo Scientific, *m/z* 622.1978) located within the source was used as a lock mass internal calibrant. Samples were introduced by an Agilent 1290 UPLC using a 10 µl injection volume for ASM samples and a 5 µl injection volume for PGM samples. Extracts were separated using a Phenomenex Kinetex 2.6 µm C18 column (2.1 mm x 50 mm) using a 9 minute, linear water-ACN gradient (from 98:2 to 2:98% water:ACN) containing 0.1% FA at a flow rate of 0.5 mL/min. The mass spectrometer was operated in data dependent positive ion mode, automatically switching between full scan MS and MS/MS acquisitions. Full scan MS spectra (*m/z* 50 - 1500) were acquired in the TOF and the top five most intense ions in a particular scan were fragmented via collision induced dissociation (CID) using the stepping function in the collision cell. LC-MS/MS data for PA mix were acquired under identical conditions. Bruker Daltonics CompassXport was used to apply lock mass calibration and convert the LC-MS/MS data from .d format to .mzXML format.

### Feature based molecular networking (FBMN) of P. aeruginosa specialized metabolism in LB and ASM formulations

MZmine (version 2.53)^133^ was used to perform feature finding on the .mzXML files associated with this experiment. The output was a feature table containing area under the curve (AUC) integration values for each feature (*m/z*-RT pair) and an .mgf file containing MS/MS spectra for each feature. The feature table was corrected for sample dilutions prior to submission to GNPS. The Feature Networking workflow (version release 22)^44^ was applied to the dilution-corrected feature table, .mgf file, and a metadata file aligning to GNPS specifications using the GNPS web platform^43^. Briefly, the data was filtered by removing all MS/MS fragment ions ± 17 Da of the precursor *m/z*. MS/MS spectra were window filtered by choosing only the top 6 fragment ions in each ± 50 Da window throughout the spectrum. The precursor ion mass tolerance was set to 0.05 Da and the MS/MS fragment ion tolerance was set to 0.1 Da. A network of nodes connected by edges was then created. Edges were filtered to have a cosine score above 0.75. Further, edges between two nodes were kept in the network only if each of the nodes appeared in each other’s respective top 5 most similar nodes. Finally, the maximum size of a molecular family was set to 50, and the lowest scoring edges were removed from molecular families until the molecular family was below this threshold. The spectra in the network were then searched against the GNPS spectral libraries. The library spectra were filtered in the same manner as the input data. All matches kept between the network spectra and library spectra were required to have a cosine score above 0.6 and at least 5 matched peaks. Sum normalization and mean aggregation were applied. The molecular network (https://gnps.ucsd.edu/ProteoSAFe/status.jsp?task=baba5df5ab1b437980f81b4e64bcffc0) was visualized using Cytoscape (version 3.7.1)^134^.

### FBMN and targeted quantification of *P. aeruginosa* specialized metabolites in SCFM1 and PGM-SCFM1 cultures

MZmine (version 2.53)^133^ was used to perform feature finding on the .mzXML files associated with this experiment. The output was a feature table containing area under the curve (AUC) integration values for each feature (*m/z*-RT pair) and an .mgf file containing MS/MS spectra for each feature. The feature table was corrected for sample dilutions prior to submission to GNPS. The Feature Networking workflow (version release 28.1)^44^ was applied to the dilution-corrected feature table, .mgf file, and a metadata file aligning to GNPS specifications using the GNPS web platform^43^. Briefly, the data was filtered by removing all MS/MS fragment ions ± 17 Da of the precursor *m/z*. MS/MS spectra were window filtered by choosing only the top 4 fragment ions in each ± 50 Da window throughout the spectrum. The precursor ion mass tolerance was set to 0.02 Da and the MS/MS fragment ion tolerance was set to 0.02 Da. A network of nodes connected by edges was then created. Edges were filtered to have a cosine score above 0.7. Further, edges between two nodes were kept in the network only if each of the nodes appeared in each other’s respective top 10 most similar nodes. Finally, the maximum size of a molecular family was set to 100, and the lowest scoring edges were removed from molecular families until the molecular family was below this threshold. The spectra in the network were then searched against the GNPS spectral libraries. The library spectra were filtered in the same manner as the input data. All matches kept between the network spectra and library spectra were required to have a cosine score above 0.7 and at least 4 matched peaks. Sum normalization and mean aggregation were applied. The molecular network (https://gnps.ucsd.edu/ProteoSAFe/status.jsp?task=5624bf8b3174483a89cf733d08ce0fff) was visualized using Cytoscape (version 3.7.1)^134^.

### Feature annotation of *P. aeruginosa* specialized metabolites

Annotations of features corresponding to commercial standards (level 1 annotation^135^) were confirmed by comparing the experimental data (exact mass, MS/MS, and retention time) with data acquired for the compound in PA mix using Bruker Daltonics DataAnalysis v4.1 (Build 362.7). Putative annotation of specialized metabolites based on matches to the GNPS libraries (level 2 annotation^135^) and molecular families (level 3 annotation^135^) were validated by comparing the experimental data (exact mass, MS/MS) to reported data^58^ and putative structures, respectively^58^. Annotated features corresponding to *P. aeruginosa* specialized metabolites identified from FBMN analysis of LB and ASM samples and SCFM1 vs PGM-SCFM1 are listed in **Table S1. PA in ASM metabolite annotation and mass defect** and **Table S2. PA in SCFM1/PGM-SCFM1 metabolite annotation and mass defect**, respectively. Multiple features (different adducts, in-source fragments, and/or in-source dimers) can represent the same metabolite.

Therefore, sum normalized AUC values for features corresponding to the same metabolite of interest (identified by co-elution and mass defect from predicted exact mass of the compound adduct, fragment, and/or dimer) were summed to provide the total AUC for a metabolite. Sum normalized AUC values for metabolites representing the same sub-structure class based upon MS/MS fragmentation pattern (e.g. HHQ-type AQs or C10-C10 containing RHLs) were summed to provide the total AUC for a sub-structure class. Sum normalized AUC values for all identified metabolites representing the same molecular family based upon MS/MS fragmentation patterns (e.g. AQs or RHLs) were summed to provide the total AUC for a molecular family.

### ICP-MS analysis of PGM

Metals were quantified from 2.5% (w/v) PGM suspended in 1X MOPS using inductively coupled plasma mass spectrometry (ICP-MS) analysis at the University of Nebraska - Lincoln Spectroscopy and Biophysics Core facility. Samples were diluted 1:1 with nitric acid, digested overnight, and diluted an additional 10-fold before conducting ICP-MS (Agilent 7500 cx). Gallium (Ga), 50 ppb, was added to each sample as an internal standard. Samples were analyzed in technical triplicate with a blank wash run between each sample. Only concentrations above the limit of quantification were retained. To determine the amount of iron added to media by the addition of PGM, or 1X MOPS when used in place of PGM, the concentration of PGM was brought to that present in most ASM formulations (0.5% w/v) and converted to micromolar concentration.

### Classical molecular networking of PGM lipophilic extracts

A classical molecular network was created from the .mzXML files using the Molecular Networking workflow (version release 27) of the GNPS web platform (gnps.ucsd.edu, https://gnps.ucsd.edu/ProteoSAFe/status.jsp?task=fc3ee81541aa4a60af38b15f08789620)^43,67^. Briefly, the data was filtered by removing all MS/MS fragment ions ± 17 Da of the precursor *m/z*. MS/MS spectra were window filtered by choosing only the top 6 fragment ions in each ± 50 Da window throughout the spectrum. The precursor ion mass tolerance was set to 0.05 Da and the MS/MS fragment ion tolerance to 0.5 Da. A network was then created where edges were filtered to have a cosine score above 0.7 and at least 4 matched fragment peaks between corresponding nodes. Further, edges between two nodes were kept in the network only if each of the nodes appeared in each other’s respective top 10 most similar nodes. Finally, the maximum size of a molecular family was set to 100, and the lowest scoring edges were removed from molecular families until the molecular family was below this threshold. The spectra in the network were then searched against the GNPS spectral libraries. The library spectra were filtered in the same manner as the input data. All matches kept between the network spectra and library spectra were required to have a cosine score above 0.6 and at least 4 matched peaks. The molecular network was visualized using Cytoscape (version 3.7.1)^134^.

### Statistical analysis

Statistical comparison of FBMN features of *P. aeruginosa* cultured in ASM formulations was conducted on the sum normalized feature table using ANOVA with Tukey’s correction for multiple comparisons in MetaboAnalyst 4.0^136–139^. Statistical comparison of individual metabolites, sub-structure classes, and molecular families performed using ANOVA with Tukey’s multiple comparisons correction in GraphPad Prism (version 8.4.0). For all analyses, *P* < 0.05 were considered statistically significant. All statistical results are summarized in Dataset S2 ASM statistical analyses.

### Data availability

All mass spectrometry data including the raw files, .mzXML files, metadata tables, MZmine settings, and Cytoscape files are available via MassIVE: MSV000086721 (LB and ASM formulations – media control, *P. aeruginosa* culture, QC, and PA mix data; corresponds to Figures 2-6); MSV000086723 (SCFM1 and PGM-SCFM1 *–* media control, *P. aeruginosa* culture, QC, and PA mix data; corresponds to Figure 7); and MSV000087153 (PGM lipophilic extracts; corresponds to Figure 8).

## ACKNOWLEDGEMENTS

We are very grateful to Mingxun Wang (University of California, San Diego) for providing input and beta versions of the networking software for metabolomics analysis; Laura Sanchez, Jacob Porter, and Katherine Zink (University of Illinois, Chicago) for providing initial collection of metabolomics data; and members of the Phelan lab for thoughtful discussion about and suggestions on improving this manuscript.

