## Supplemental Material for "Commercial porcine gastric mucin contributes to variation in production of small molecule virulence factors by *Pseudomonas aeruginosa* when cultured in different formulations of artificial sputum medium"

**Supplemental Material Table of Contents**

Table S1 PA in ASM metabolite annotation and mass defect…………………………….................. 3

Table S2 PA in SCFM1/PGM-SCFM1 metabolite annotation and mass defect............................... 4

Reference............................................................................................................................ 5

#### Table S1 PA in ASM metabolite annotation and mass defect.

| Molecular Family | ID | Annotation | Adduct | Measured *m/z* | Calculated *m/z* | Mass Defect (ppm) | Annotation Level[#] |
|---|---|---|---|---|---|---|---|
| Phenazine | 10 | 1-HP | [M+H] | 197.0702 | 197.0709 | -3.55 | 1 |
| Phenazine | 16 | PYO | [M+H] | 211.0858 | 211.0866 | -3.79 | 1 |
| Phenazine | 226 | PYO | [M+H] | 211.086 | 211.0866 | -2.84 | 1 |
| Phenazine | 34 | PCN | [M+H] | 224.0813 | 224.0818 | -2.23 | 1 |
| Phenazine | 8 | PCA | [M+H] | 225.0651 | 225.0659 | -3.55 | 1 |
| Quinolone | 76 | C5-HQ | [M+H] | 216.1377 | 216.1383 | -2.78 | 2 |
| Quinolone | 391 | C6-HQ | [M+H] | 230.1535 | 230.1539 | -1.74 | 2 |
| Quinolone | 72 | C5-QNO | [M+H] | 232.1325 | 232.1332 | -3.02 | 2 |
| Quinolone | 1 | C7-HQ (HHQ) | [M+H] | 244.1688 | 244.1696 | -3.28 | 1 |
| Quinolone | 189 | C6-QNO | [M+H] | 246.1482 | 246.1489 | -2.84 | 2 |
| Quinolone | 117 | C8:1-HQ | [M+H] | 256.169 | 256.1696 | -2.34 | 2 |
| Quinolone | 51 | C7:1-HQ-OH | [M+H] | 258.1482 | 258.1489 | -2.71 | 3 |
| Quinolone | 32 | C8-HQ | [M+H] | 258.1845 | 258.1852 | -2.71 | 2 |
| Quinolone | 3 | C7-QNO (HQNO) | [M+H] | 260.1638 | 260.1645 | -2.69 | 1 |
| Quinolone | 4 | PQS | [M+H] | 260.1638 | 260.1645 | -2.69 | 1 |
| Quinolone | 5 | C9:1-HQ | [M+H] | 270.1845 | 270.1852 | -2.59 | 2 |
| Quinolone | 436 | C9:1-HQ | [M+H] | 270.1847 | 270.1852 | -1.85 | 2 |
| Quinolone | 2 | C9-HQ (NHQ) | [M+H] | 272.2001 | 272.2009 | -2.94 | 2 |
| Quinolone | 48 | C8-QNO | [M+H] | 274.1795 | 274.1802 | -2.55 | 2 |
| Quinolone | 399 | C9:2-PQS | [M+H] | 284.1633 | 284.1645 | -4.22 | 3 |
| Quinolone | 105 | C10:1-HQ | [M+H] | 284.2003 | 284.2009 | -2.11 | 2 |
| Quinolone | 369 | C10:1-HQ | [M+H] | 284.2005 | 284.2009 | -1.41 | 2 |
| Quinolone | 179 | C9:1-PQS | [M+H] | 286.1795 | 286.1802 | -2.45 | 3 |
| Quinolone | 11 | C9:1-QNO<br>C9:1-PQS | [M+H] | 286.1795 | 286.1802 | -2.45 | 2 |
| Quinolone | 291 | C9:1-HQ-OH | [M+H] | 286.1795 | 286.1802 | -2.45 | 3 |
| Quinolone | 7 | C9-QNO (NQNO) | [M+H] | 288.1951 | 288.1958 | -2.43 | 2 |
| Quinolone | 44 | C9-PQS | [M+H] | 288.1951 | 288.1958 | -2.43 | 2 |
| Quinolone | 77 | C9-HQ-OH | [M+H] | 288.1953 | 288.1958 | -1.73 | 3 |
| Quinolone | 205 | C11:2-HQ | [M+H] | 296.1995 | 296.2009 | -4.73 | 3 |
| Quinolone | 9 | C11:2-HQ | [M+H] | 296.2004 | 296.2009 | -1.69 | 3 |
| Quinolone | 225 | C11:1-HQ | [M+H] | 298.2156 | 298.2165 | -3.02 | 2 |
| Quinolone | 236 | C11:1-HQ | [M+H] | 298.2157 | 298.2165 | -2.68 | 2 |
| Quinolone | 307 | C10:1-QNO<br>C10:1-PQS | [M+H] | 300.1955 | 300.1958 | -1.00 | 2 |
| Quinolone | 19 | C11-HQ | [M+H] | 300.2315 | 300.2322 | -2.33 | 2 |
| Quinolone | 324 | C9-QNO-OH<br>C9-PQS-OH | [M+H] | 304.1901 | 304.1907 | -1.97 | 3 |
| Quinolone | 288 | C12:1-HQ | [M+H] | 312.2318 | 312.2322 | -1.28 | 2 |
| Quinolone | 31 | C11:1-QNO<br>C11:1-PQS | [M+H] | 314.2109 | 314.2115 | -1.91 | 2 |
| Quinolone | 83 | C11-QNO | [M+H] | 316.2265 | 316.2271 | -1.90 | 2 |
| Quinolone | 388 | C13:2-HQ | [M+H] | 324.2317 | 324.2322 | -1.54 | 3 |

| | | | | | | | |
|---|---|---|---|---|---|---|---|
| Quinolone | 17 | C13:2-HQ | [M+H] | 324.2317 | 324.2322 | -1.54 | 3 |
| Quinolone | 63 | C13:1-HQ | [M+H] | 326.2473 | 326.2478 | -1.53 | 2 |
| Quinolone | 172 | C13-HQ | [M+H] | 328.2629 | 328.2635 | -1.83 | 2 |
| Quinolone | 67 | C13:2-QNO<br>C13:2-PQS | [M+H] | 340.2269 | 340.2271 | -0.59 | 3 |
| Quinolone | 283 | C13:1-QNO<br>C13:1-PQS | [M+H] | 342.2422 | 342.2428 | -1.75 | 3 |
| Quinolone | 160 | C15:2-HQ | [M+H] | 352.2627 | 352.2635 | -2.27 | 3 |
| Quinolone | 315 | C15:1-HQ | [M+H] | 354.2780 | 354.2791 | -3.10 | 3 |
| Quinolone | 106 | C17:1-HQ | [M+H] | 382.3088 | 382.3104 | -4.18 | 3 |
| Rhamnolipid | 149 | Rha-C10-C10 | [M+H] | 505.3371 | 505.3371 | 0 | 1 |
| Rhamnolipid | 30 | Rha-C10-C10 | [M+Na] | 527.3188 | 527.3191 | -0.57 | 1 |
| Rhamnolipid | 124 | Rha-C10-C12:1 | [M+Na] | 553.3347 | 553.3347 | 0 | 1 |
| Rhamnolipid | 451 | Rha-C10-C12 | [M+H] | 533.3665 | 533.3684 | -3.56 | 1 |
| Rhamnolipid | 139 | Rha-C10-C12 | [M+Na] | 555.3504 | 555.3504 | 0 | 1 |
| Rhamnolipid | 40 | Rha-Rha-C10-C10 | [M+H] | 651.3948 | 651.3950 | -0.31 | 1 |
| Rhamnolipid | 133 | Rha-Rha-C10-C10 | [2M+K+H] | 670.3689 | 670.3730 | -6.12 | 1 |
| Rhamnolipid | 45 | Rha-Rha-C10-C10 | [M+Na] | 673.3767 | 673.3770 | -0.45 | 1 |
| Rhamnolipid | 154 | Rha-Rha-C10-C12:1 | [M+H] | 677.4109 | 677.4107 | 0.30 | 1 |
| Rhamnolipid | 110 | Rha-Rha-C10-C12:1 | [M+Na] | 699.3928 | 699.3926 | 0.29 | 1 |
| Rhamnolipid | 88 | Rha-Rha-C10-C12 | [M+H] | 679.4263 | 679.4263 | 0 | 1 |
| Rhamnolipid | 71 | Rha-Rha-C10-C12 | [M+Na] | 701.408 | 701.4083 | -0.43 | 1 |
| Siderophore | 300 | PCH | [M+H] | 325.0673 | 325.0675 | -0.62 | 2 |
| Siderophore | 298 | PCH | [M+H] | 325.0673 | 325.0675 | -0.62 | 2 |
| Siderophore | 426 | PVD E | [M+2H] | 667.3094 | 667.3102 | -1.20 | 2 |
| Siderophore | 428 | PVD E | [M+3H] | 445.2085 | 445.2092 | -1.57 | 2 |
| Siderophore | 410 | PVD E + Fe | [M+2H] | 693.7660 | 693.7660 | 0 | 2 |
| Siderophore | 411 | PVD E + Fe | [M+3H] | 462.8470 | 462.8464 | 1.30 | 2 |
| Siderophore | 424 | Ferribactin | [M+2H] | 676.3333 | 676.3337 | -0.59 | 3 |
| Siderophore | 378 | Ferribactin | [M+3H] | 451.2249 | 451.2249 | 0 | 3 |

[#] Annotation level according to [1]

**Table S2 PA in SCFM1/PGM-SCFM1 metabolite annotation and mass defect.**

| Molecular Family | ID | Annotation | Adduct | Measured *m/z* | Calculated *m/z* | Mass Defect (ppm) | Annotation Level# |
|---|---|---|---|---|---|---|---|
| Phenazine | 32 | 1-HP | [M+H] | 197.0705 | 197.0709 | -2.03 | 1 |
| Phenazine | 13 | PYO | [M+H] | 211.0859 | 211.0866 | -3.32 | 1 |
| Phenazine | 26 | PCA | [M+H] | 225.0653 | 225.0659 | -2.67 | 1 |
| Quinolone | 1 | C7-HQ (HHQ) | [M+H] | 244.1691 | 244.1696 | -2.05 | 1 |
| Quinolone | 3 | C7-QNO (HQNO) | [M+H] | 260.1641 | 260.1645 | -1.54 | 1 |
| Quinolone | 124 | C9:1-HQ | [M+H] | 270.1846 | 270.1852 | -2.22 | 2 |
| Quinolone | 7 | C9:1-HQ | [M+H] | 270.1847 | 270.1852 | -1.85 | 2 |
| Quinolone | 2 | C9-HQ (NHQ) | [M+H] | 272.2004 | 272.2009 | -1.84 | 2 |
| Quinolone | 504 | C9:1-QNO/<br>C9:1-PQS | [M+H] | 286.1794 | 286.1802 | -2.8 | 2 |
| Quinolone | 24 | C9:1-HQ-OH | [M+H] | 286.1797 | 286.1802 | -1.75 | 2 |
| Quinolone | 4 | C9-QNO (NQNO) | [M+H] | 288.1955 | 288.1958 | -1.04 | 2 |
| Rhamnolipid | 313 | Rha-C10-C10 | [M+H] | 505.3358 | 505.3371 | -2.57 | 1 |
| Rhamnolipid | 90 | Rha-C10-C10 | [M+Na] | 527.3188 | 527.3191 | -0.57 | 1 |
| Rhamnolipid | 343 | Rha-C10-C12 | [M+Na] | 555.3493 | 555.3504 | -1.98 | 1 |
| Rhamnolipid | 61 | Rha-Rha-C10-C10 | [M+H] | 651.3933 | 651.3950 | -2.61 | 1 |
| Rhamnolipid | 47 | Rha-Rha-C10-C10 | [M+Na] | 673.3759 | 673.3770 | -1.63 | 1 |
| Rhamnolipid | 417 | Rha-Rha-C10-C12:1 | [M+H] | 677.4094 | 677.4107 | -1.92 | 1 |
| Rhamnolipid | 269 | Rha-Rha-C10-C12:1 | [M+Na] | 699.3916 | 699.3926 | -1.43 | 1 |
| Rhamnolipid | 176 | Rha-Rha-C10-C12 | [M+H] | 679.4255 | 679.4263 | -1.18 | 1 |
| Rhamnolipid | 102 | Rha-Rha-C10-C12 | [M+Na] | 701.407 | 701.4083 | -1.85 | 1 |
| Siderophore | 15 | PCH | [M+H] | 325.0669 | 325.0675 | -1.85 | 2 |
| Siderophore | 8 | PCH | [M+H] | 325.067 | 325.0675 | -1.54 | 2 |
| Siderophore | 122 | PVD E | [M+2H] | 667.3084 | 667.3102 | -2.7 | 2 |
| Siderophore | 506 | Ferribactin | [M+2H] | 676.3315 | 676.3337 | -3.25 | 3 |
| Siderophore | 609 | Ferribactin | [M+3H] | 451.2237 | 451.2249 | -2.66 | 3 |

### Annotation level according to [1]
